# A bioinformatics approach to systematically analyze the molecular patterns of monkeypox virus-host cell interactions

**DOI:** 10.1101/2022.10.12.511850

**Authors:** Zhongxiang Tang, Yu Mao, Yuting Meng, Xiangjie Qiu, Ousman Bajinka, Guojun Wu, Yurong Tan

## Abstract

Monkeypox has been spreading worldwide since May 2022, when the World Health Organization (WHO) declared the outbreak of a “Public health emergency of international concern”. The spread of monkeypox has posed a serious threat to the health of people all over the world but few studies have been carried out on it, and the molecular mechanism of monkeypox after infection remains unclear. We therefore implemented a transcriptome analysis to identify signaling pathways and biomarkers in monkeypox-infected cells to help understand monkeypox-host cell interactions. In this study, the dataset GSE36854 and GSE11234 were obtained from GEO. Among them, 84 significantly different genes were identified in the dataset GSE36854, followed by KEGG, GO analysis protein-protein interaction (PPI) construction and Hub gene extraction. We also analyzed the expression regulation patterns of hub genes and screened the drugs targeting the hub genes. The results showed that monkeypox-infected cells significantly activated the cellular immune response, and induced inflammatory response. IER3, IFIT2, IL11, ZC3H12A, EREG, IER2, NFKBIE, FST, IFIT1 and AREG were the top 10 hub genes, in which anti-viral gene IFIT1 and IFIT2 were significantly suppressed. AP-26113 and itraconazole promoting the expression of IFIT1 and IFIT2 may be used as new candidates for the treatment of monkeypox viral infection. Our results provide a new entry point for understanding the mode of interaction between monkeypox virus and its host.

## Introduction

Monkeypox is a viral zoonosis caused by monkeypox virus infection and used to occur mostly in West and Central Africa[1, 2]. However, since May 2022, many countries and regions outside Africa have reported cases of monkeypox. There is no direct epidemiological link between this outbreak and the Central and West African regions. The WHO declared the multinational monkeypox outbreak a “Public health emergency of international concern” on July 23^rd^ 2022[3]. As of 22rd September, 2022, more than 64,000 confirmed cases of monkeypox and 135 deaths have been reported in 107 countries and areas affected by the outbreak.

Monkeypox, like smallpox and vaccinia viruses is a member of the orthopoxvirus family and is usually less severe than smallpox[4]. In general, Monkeypox is a self-limiting disease with symptoms lasting 2 to 4 weeks and a mortality rate of about 3.6%.; with a mortality of 3.6% in the West African branch and 10.6% in the Congo Basin branch[5]. Incubation period of Monkeypox is usually 5 to 21 days, which is not infectious. The prodromal symptoms usually include fever, headache, lymphadenopathy, muscle soreness and fatigue. Among them, lymphadenopathy is a prominent feature of monkeypox and can be distinguished from similar diseases such as chickenpox, measles and smallpox[6]. Rashes usually appear within 3 days of onset, first on the face and then on the limbs (hands, legs, and feet). Oral mucosa, genitals, conjunctiva, and cornea may also be present. Animal-to-human transmission is the main route of transmission through animal bite or direct contact with animal blood, body fluids, skin or mucosal lesions, etc., or inappropriately cooked animals infected with monkeypox virus. Human-to-human transmission includes respiratory droplets, direct contact or sexual transmission[7, 8]. The current evidence for suspected mother-to-child is insufficient[9]. Some of the people affected by this outbreak are MSM (Men who have sex with men)[10]. Currently, researchers have also reported cases of monkeypox in human-to-animal transmission[11], but there are also cases of monkeypox combined with Sars-Cov-2 infection[12], which require an urgent need to the study of monkeypox.

In view of the severity of the monkeypox epidemic, in addition to the need to clarify the many reasons for the outbreak and find ways to limit the spread of monkeypox, there is also an urgent need to elucidate the mechanisms by which monkeypox interacts with humans to cause severe disease. Moreover, to find a treatment for monkeypox. In this study, we obtained the sequence data of the transcripts of human cells infected by monkeypox in GEO and screened the significantly different genes, analyzed the signal pathway, expression regulation and metabolic pathway involved in the significantly different genes. We also compared the difference between monkeypox, cowpox and vaccinia in the change of human cell transcripts after infection. Furthermore, the interaction between monkeypox and human cells was elucidated to find a target for the treatment of monkeypox.

## Methods

### Acquisition of sequencing data and analysis of differential genes

To determine the effect of monkeypox infection on human cells at the transcript level, we downloaded the sequencing datasets GSE36854 and GSE11234 from the GEO Database[13]. The data set GSE36854 was sequenced on Agilent’s GPL4133. The data set contained eight samples; two of which were Vaccinia virus strain IHD-W, two were Vaccinia virus strain Brighton Red or Monkeypox virus strain MSF # 6 infected Hela cells, two were blank controls. The data set GSE11234 is based on GPL6763, which contains many types of poxvirus and human genome template information. We extracted only samples of monkeypox-infected Hela cells for analysis of the genomic alterations. We processed and visualized the sequencing data using R (R version 4.1.3) language in which the Limma package was used to screen for significantly differentially expressed genes[14]. |LogFC| > 1 and | P <0.05 | was used as screening conditions.

### Pathway enrichment and Gene ontology analysis

Pathway enrichment and Gene ontology analysis for significantly different genes can effectively establish the intrinsic association among multiple genes, this in turn helps us understand cell signaling, gene regulatory networks, and the composition of biological systems. GO analysis Gene Ontology can be divided into three parts: Molecular Function, biological process and cellular component, which can systematically describe the function, biological process and cellular localization of significantly different genes. We performed KEGG and GO analysis using the online analysis platform enrichr (enrichr database is a comprehensive online tool for geneenrichment analysis that contains a large number of genome annotation libraries for analysis. Moreover, we download Transcription, PATHWAYS, Ontologies, Diseases/Drugs, Cell types, etc.)[15]. KEGG 2021 was used as the database of pathway enrichment analysis, and the pathway differences of significantly different genes were compared among the three virus-infected Hela cells. P < 0.05 was used as the criterion of significance. GSEA analysis was performed to identify pathways based on the obtained transcript sequences of monkeypox infection[16].

### Identification of transcription factors and miRNAs that regulate DEGs

The mode of expression regulation of significantly different genes is the underlying logic of the significant changes in the intrinsic mode of cells infected with monkeypox. Therefore, elucidating the expression regulation of significantly different genes will help us to find the key to the occurrence of monkeypox infection. It will also help us to target key cell-intrinsic gene links involved in the pathophysiological response from monkeypox infection. We analyzed significant differences in monkeypox-infected Hela cells using the online analysis tool NetworkAnalyst 3.0 (https://www.networkanalyst.ca/) (for gene expression analysis and meta-analysis)[17]. The ENCODE database (transcription factor and gene target data derived from the ENCODE ChIP-seq data) was used to analyze the expression regulation relationship between miRNAs and DEGs [18]. The TarBase V8.0 database (comprehensive experimentally validated miRNA-gene interaction data collected from TarBase) was used to analyze the expression regulation relationship between miRNAs and DEGs[19, 20].

### Protein-protein interaction network analysis and hub gene extraction

We constructed the PPI network using the String (https://string-db.org, version: 11.5) database of significantly different genes for monkeypox infection to explore possible relationships among DEGs[21]. These include, but are not limited to, co-expression, and physical relationship. We also used the Density of Maximum Neighborhood Component (DMNC) method in the Cytohubba plug-in in Cytoscape (version:3.8.2) to extract the top 10 genes in this PPI as hub genes[22, 23].

### ENCODE data verify the binding of transcription factors

The Encode (https://www.encodeproject.org/) database was used to analyze the binding of IRF1to the IFITs promoter and visualized with the WashU Epigenome Browser online tool (http://epigenomegateway.wustl.edu/browser/)[18, 24].

### Drug sensitivity and gene-disease association analysis

We used the Gene-disease Associations tool of network analysts in the analysis platform to analyze the diseases involved by significantly different genes based on the DisGeNET database(The literature curated gene-disease association information was collected from the DisGeNET database [25]. The expression data of IFIT1 and IFIT2 in the NCI-60 cell line drug database was downloaded from the CellMiner database. The Pearson correlation coefficient with 792 FDA-approved drugs in clinical trials was analyzed, and the results were screened and visualized (p < 0.05)[26].

## Results

### Screening of differentially expressed genes

We first compared the expression profiles of monkeypox virus-infected and mock-infected Hela cells in the dataset GSE36854 to determine the regulation of cellular transcription by monkeypox infection.We screened 126 significant different genes, 24 of which were significantly down-regulated and 102 were significantly up-regulated (Fig. 1A). Of the 102 significantly upregulated genes, 43 were histone-coding genes, but the transcriptional upregulation of histone mRNAs was due to the experimental illusion caused by de novo polyadenylation of these mRNAs by the viral poly (a) polymerase. It was removed in subsequent analyses. In addition, we determined the transcriptional regulation of cells infected by cowpox virus and Vaccinia virus. We found that cowpox virus resulted in 390 genes significantly changed, including 168 significantly up-regulated genes and 222 significantly down-regulated genes. Vaccinia virus resulted in 390 significant gene changes, including 131 significantly up-regulated genes and 97 significantly down-regulated genes. We then compared the expression profiles of monkeypox virus genes after monkeypox infection and mock infection in the dataset GSE11234 to identify early genes expressed in Hela cells after monkeypox infection (Fig. 1B). The dataset has point-in-time information.

**Fig. 1.**
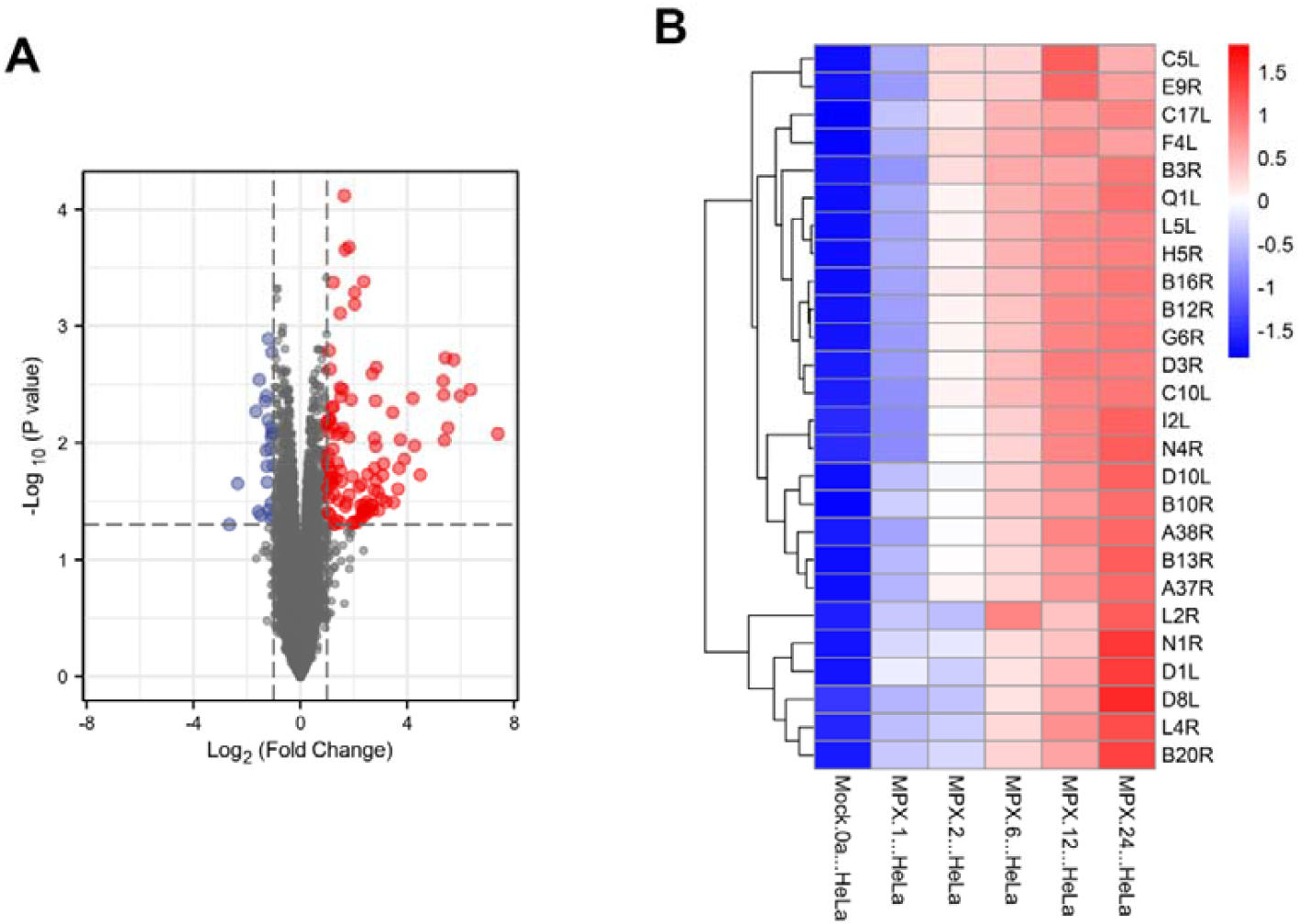
Screening of significantly different genes. A, volcano plot of differential genes in monkeypox-infected Hela cells in the GSE36854 data set. B, heat map of monkeypox early genes in monkeypox-infected Hela cells in the GSE11234 data set.

### GO and pathway enrichment analysis

The results showed that the signal pathways of monkeypox and vaccinia were mainly enriched in TNF signaling pathway, Il-17 signaling pathway and NF-kappaB signaling pathway. However vaccinia altered signaling pathways are mainly enriched in Parathyroid hormone synthesis, secretion and action, ErbB signaling pathway, colorectal cancer, Neomycin, kanamycin and gentamicin biosynthesis, carbohydrate digestion and absorption, other types of O-glycan biosynthesis (Figure 2). It is suggested that monkeypox and vaccinia may cause more severe stress and pathological state than vaccina. The GO analysis also showed that monkeypox infection activated a variety of chemotactic signaling pathways. Table 1 summarizes the top 10 terms in the class of biological processes, molecular functions, and cellular components involved in differentially expressed genes. To determine which key chemotactic responses after monkeypox infection, we analyzed changes in chemokines and MHC molecules after infection in three viruses. The results showed that CXCL1 was significantly activated after vaccinia and monkeypox infection (Figure 3B), However, MHC molecules were largely unchanged after viral infection (Figure 3A), this suggests that CXCL1 plays a key role in controlling chemotaxis after monkeypox infection.

**Table 1.**
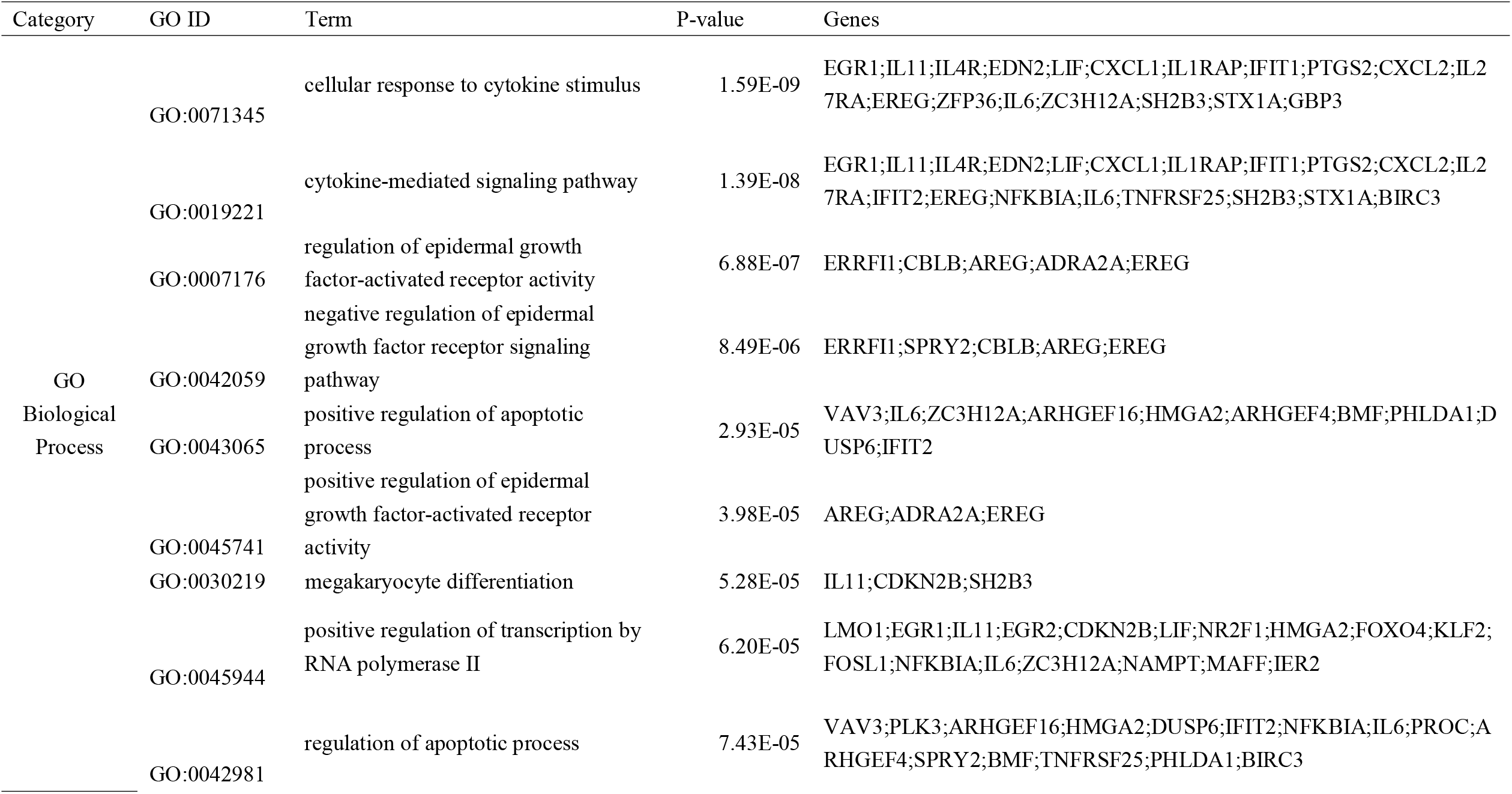
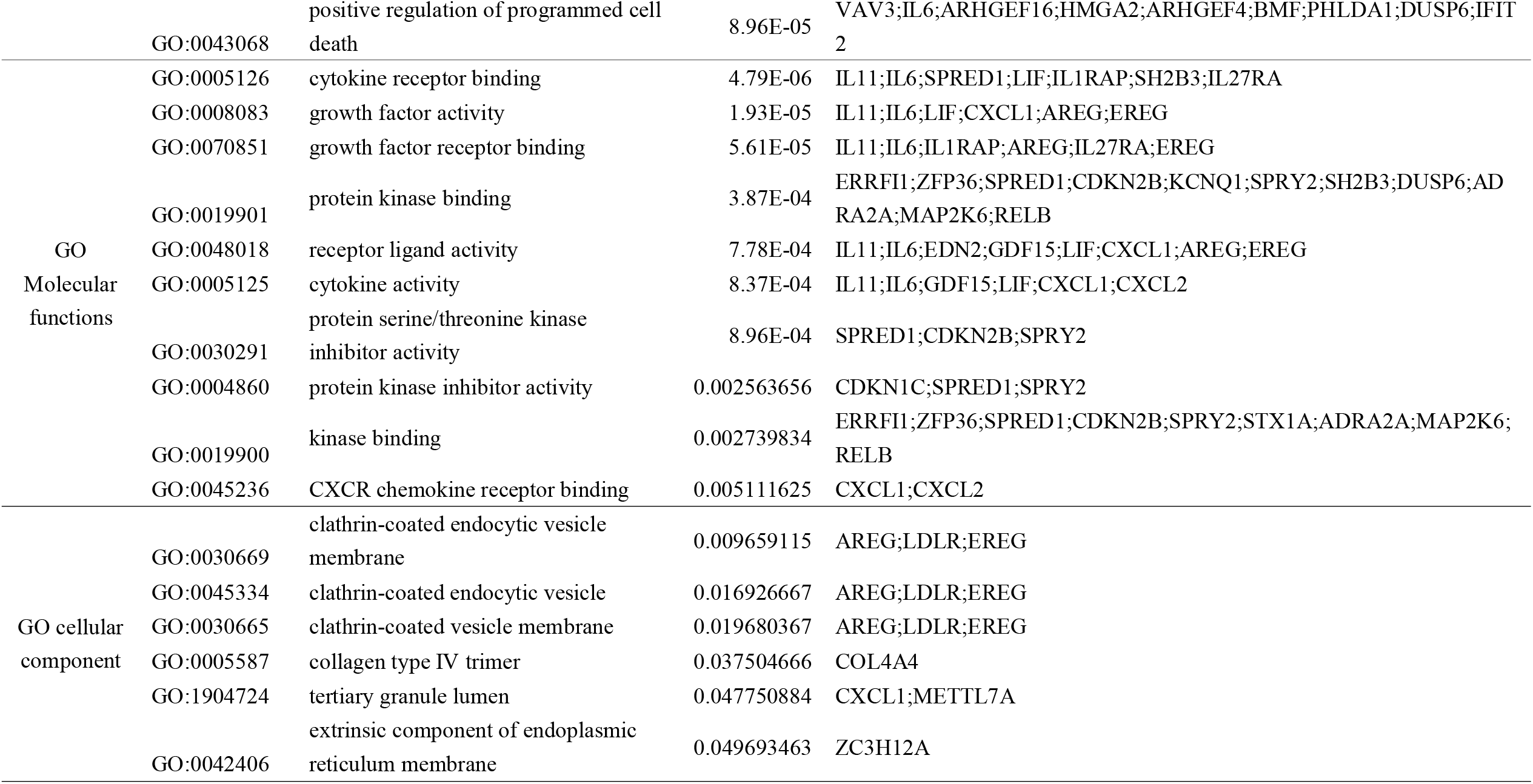
biological processes, molecular functions and cellular components involved in differentially expressed genes Category GO ID Term P-value Genes

| Category | GO ID | Term | P-value | Genes |
| --- | --- | --- | --- | --- |
| GO<br>Biological<br>Process | GO:0071345 | cellular response to cytokine stimulus | 1.59E-09 | EGR1;IL11;IL4R;EDN2;LIF;CXCL1;IL1RAP;IFIT1;PTGS2;CXCL2;IL27RA;EREG;ZFP36;IL6;ZC3H12A;SH2B3;STX1A;GBP3 |
|  | GO:0019221 | cytokine-mediated signaling pathway | 1.39E-08 | EGR1;IL11;IL4R;EDN2;LIF;CXCL1;IL1RAP;IFIT1;PTGS2;CXCL2;IL27RA;IFIT2;EREG;NFKBIA;IL6;TNFRSF25;SH2B3;STX1A;BIRC3 |
|  | GO:0007176 | regulation of epidermal growth factor-activated receptor activity | 6.88E-07 | ERRFI1;CBLB;AREG;ADRA2A;EREG |
|  | GO:0042059 | negative regulation of epidermal growth factor receptor signaling pathway | 8.49E-06 | ERRFI1;SPRY2;CBLB;AREG;EREG |
|  | GO:0043065 | positive regulation of apoptotic process | 2.93E-05 | VAV3;IL6;ZC3H12A;ARHGEF16;HMGA2;ARHGEF4;BMF;PHLDA1;DUSP6;IFIT2 |
|  | GO:0045741 | positive regulation of epidermal growth factor-activated receptor activity | 3.98E-05 | AREG;ADRA2A;EREG |
|  | GO:0030219 | megakaryocyte differentiation | 5.28E-05 | IL11;CDKN2B;SH2B3 |
|  | GO:0045944 | positive regulation of transcription by RNA polymerase II | 6.20E-05 | LMO1;EGR1;IL11;EGR2;CDKN2B;LIF;NR2F1;HMGA2;FOXO4;KLF2;FOSL1;NFKBIA;IL6;ZC3H12A;NAMPT;MAFF;IER2 |
|  | GO:0042981 | regulation of apoptotic process | 7.43E-05 | VAV3;PLK3;ARHGEF16;HMGA2;DUSP6;IFIT2;NFKBIA;IL6;PROC;ARHGEF4;SPRY2;BMF;TNFRSF25;PHLDA1;BIRC3 |
|  | GO:0043068 | positive regulation of programmed cell death | 8.96E-05 | VAV3;IL6;ARHGEF16;HMGA2;ARHGEF4;BMF;PHLDA1;DUSP6;IFIT2 |
| GO Molecular functions | GO:0005126 | cytokine receptor binding | 4.79E-06 | IL11;IL6;SPRED1;LIF;IL1RAP;SH2B3;IL27RA |
|  | GO:0008083 | growth factor activity | 1.93E-05 | IL11;IL6;LIF;CXCL1;AREG;EREG |
|  | GO:0070851 | growth factor receptor binding | 5.61E-05 | IL11;IL6;IL1RAP;AREG;IL27RA;EREG |
|  | GO:0019901 | protein kinase binding | 3.87E-04 | ERRFI1;ZFP36;SPRED1;CDKN2B;KCNQ1;SPRY2;SH2B3;DUSP6;ADRA2A;MAP2K6;RELB |
|  | GO:0048018 | receptor ligand activity | 7.78E-04 | IL11;IL6;EDN2;GDF15;LIF;CXCL1;AREG;EREG |
|  | GO:0005125 | cytokine activity | 8.37E-04 | IL11;IL6;GDF15;LIF;CXCL1;CXCL2 |
|  | GO:0030291 | protein serine/threonine kinase inhibitor activity | 8.96E-04 | SPRED1;CDKN2B;SPRY2 |
|  | GO:0004860 | protein kinase inhibitor activity | 0.002563656 | CDKN1C;SPRED1;SPRY2 |
|  | GO:0019900 | kinase binding | 0.002739834 | ERRFI1;ZFP36;SPRED1;CDKN2B;SPRY2;STX1A;ADRA2A;MAP2K6;RELB |
|  | GO:0045236 | CXCR chemokine receptor binding | 0.005111625 | CXCL1;CXCL2 |
| GO cellular component | GO:0030669 | clathrin-coated endocytic vesicle membrane | 0.009659115 | AREG;LDLR;EREG |
|  | GO:0045334 | clathrin-coated endocytic vesicle | 0.016926667 | AREG;LDLR;EREG |
|  | GO:0030665 | clathrin-coated vesicle membrane | 0.019680367 | AREG;LDLR;EREG |
|  | GO:0005587 | collagen type IV trimer | 0.037504666 | COL4A4 |
|  | GO:1904724 | tertiary granule lumen | 0.047750884 | CXCL1;METTL7A |
|  | GO:0042406 | extrinsic component of endoplasmic reticulum membrane | 0.049693463 | ZC3H12A |

**Fig. 2.**
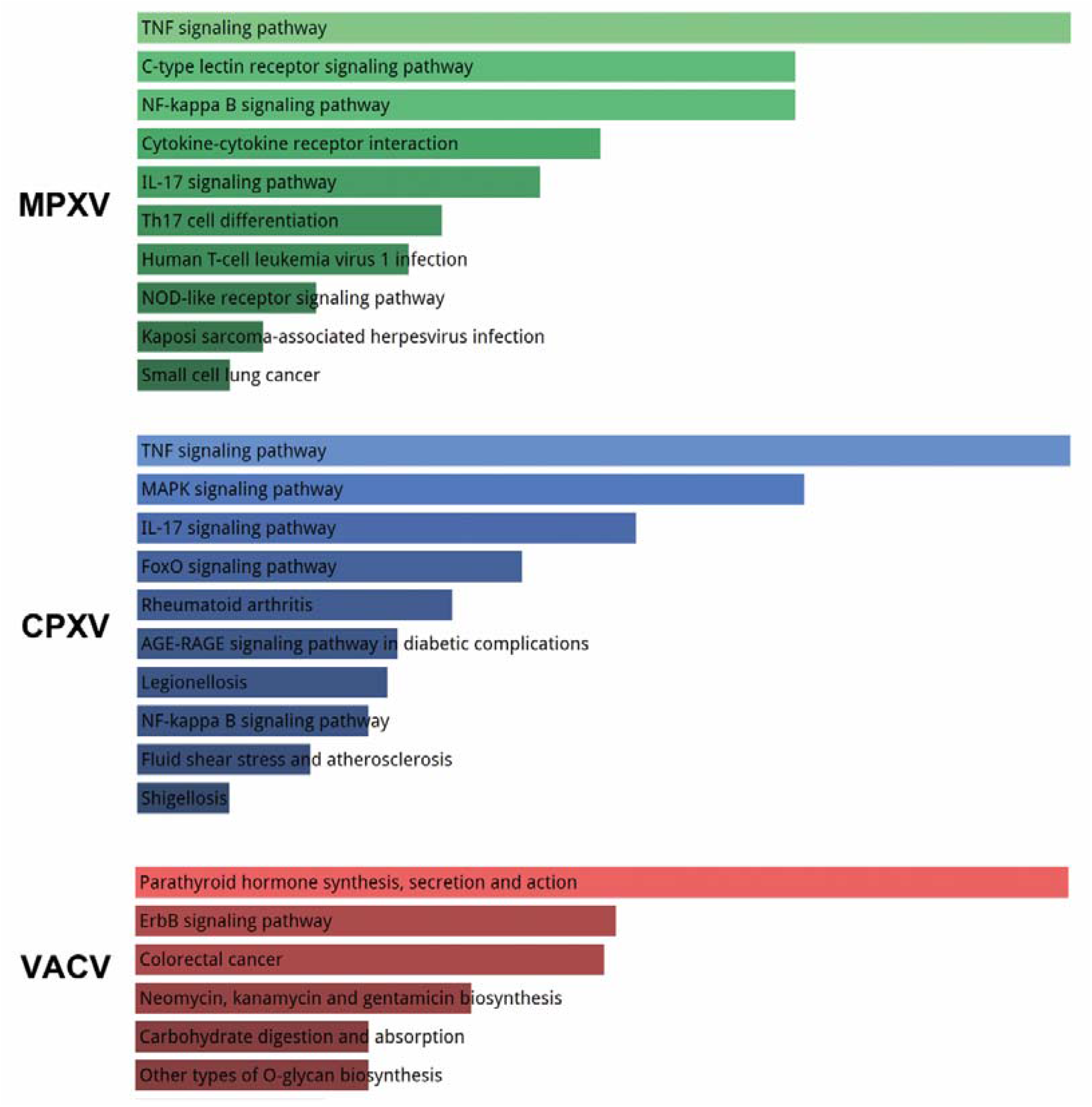
KEGG signaling pathway with significantly different gene enrichment produced by MPXV, CPCV and VACV in HELA cells.

**Fig. 3.**
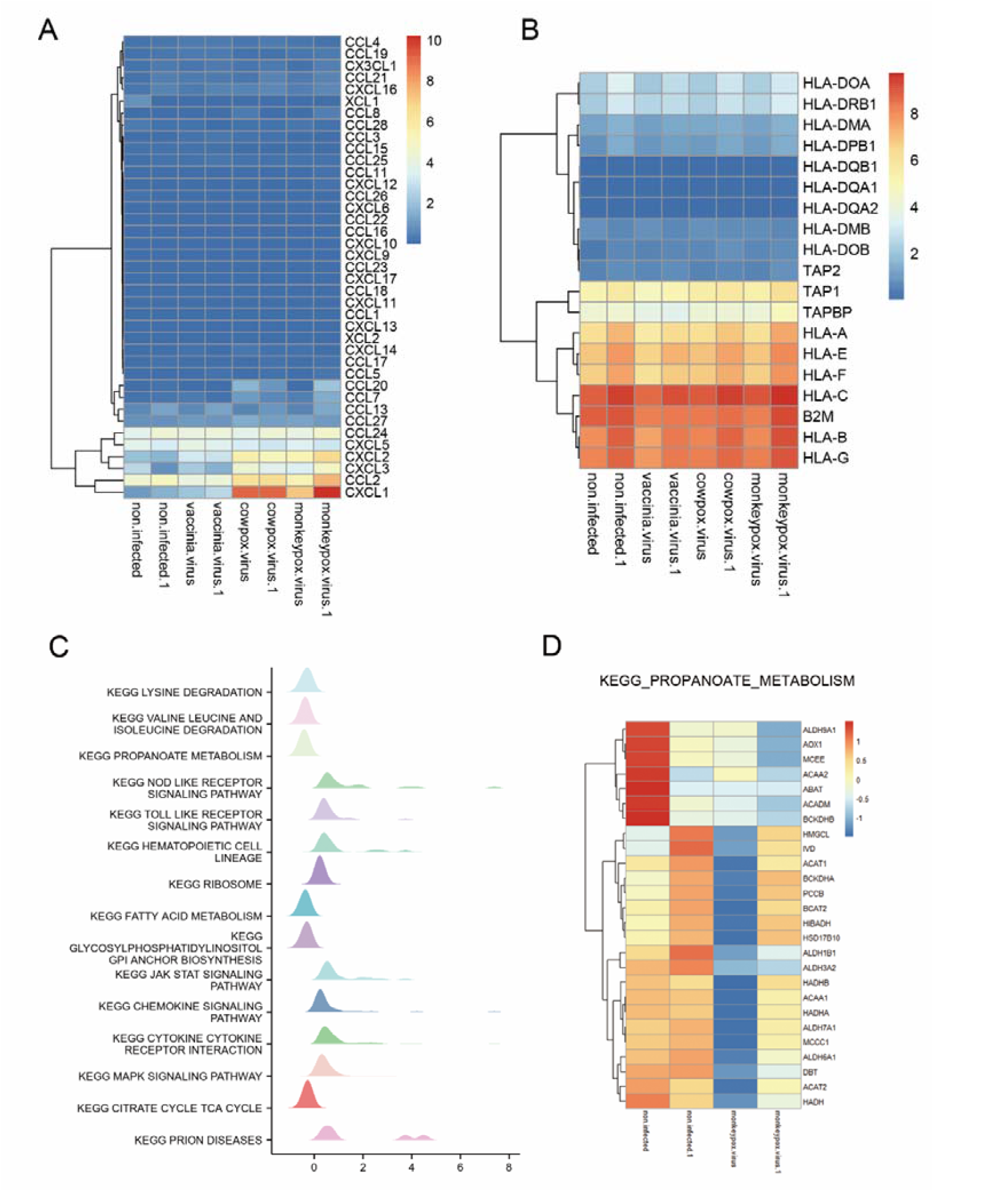
GSEA analysis of the signaling pathway in monkeypox-infected Hela cells. (A) heatmap of chemokines expression in monkeypox, vaccinia and vaccinia-infected Hela cells, (b) Heatmap of MHC molecules expression in monkeypox, vaccinia and vaccinia-infected Hela cells, (C) Gene changes in monkeypox-infected Hela cells analyzed by GSEA, (C) Gene changes in monkeypox-infected Hela cells.(D) heat map depicts the signal pathway changes in Propanoate metabolism.

### GSEA analysis

To more precisely characterize the genetic changes in cells following monkeypox infection, we performed GSEA analysis. We found that monkeypox viral infection significantly activated intracellular pattern recognition receptor pathways, such as NOD and TOLL, which is consistent with our KEGG and GO analyses of the significant different genes. But monkeypox viral infection in particular also significantly reduced metabolic levels within cells, including LYSINE_DEGRADATION, VALINE_LEUCINE_AND_ISOLEUCINE_DEGRADATION, PROPANOATE_METABOLISM,FATTY_ACID_METABOLISM, GLYCOSYLPHOSPHATIDYLINOSITOL_GPI_ANCHOR_BIOSYNTHESIS 和 CITRATE_CYCLE_TCA_CYCLE (Fig. 3C). We compared the differences of all genes of the PROPANOATE pathway before and after infection and found that monkeypox viral infection inhibited the expression of all genes of this pathway (Fig. 3D). This may be due to the hijacking of cellular metabolites after infection as a substrate for the viral replication.

### Identification of disease associations

In order to clarify the relationship between disease and the significantly altered genes in the host after monkeypox viral infection, we used the Gene-disease Associations tool of network analysts in the analysis platform to analyze the diseases involved by significantly different genes. The results showed that Liver Cirrhosis, Schizophrenia, Myocardial Ischemia, Inflammation, Reperfusion Injury, Mammary Neoplasms, Hypertensive disease, Brain Ischemia, Mental Depression and Juvenile arthritis were the 10 diseases with the highest degree of association, and IL6 and PTGS2 were the two genes with the highest degree of association with the diseases (Fig. 4).

**Fig. 4.**
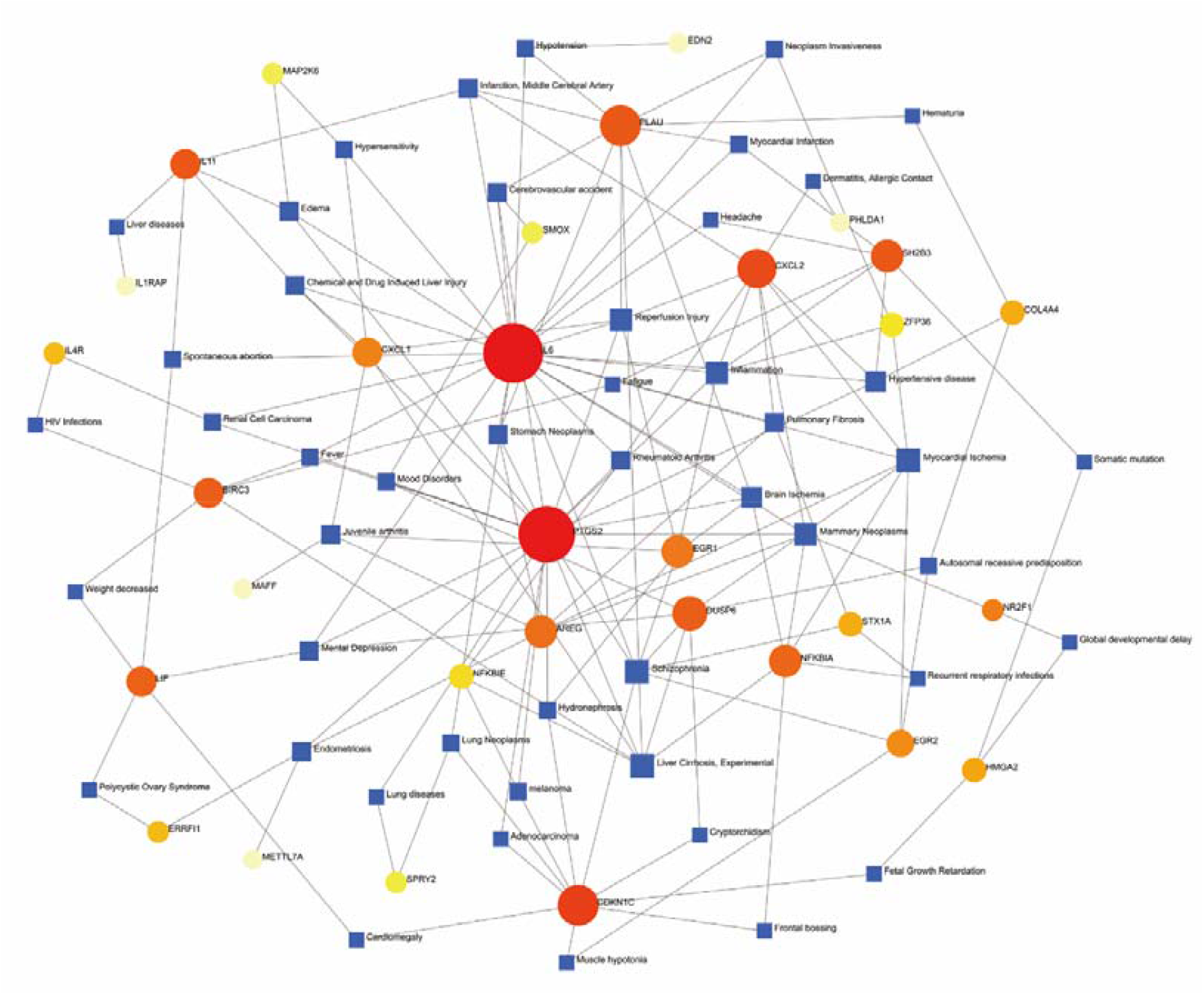
DEGs and disease-related networks. Circular nodes represent genes, and square nodes represent disease types.

**Fig. 5.**
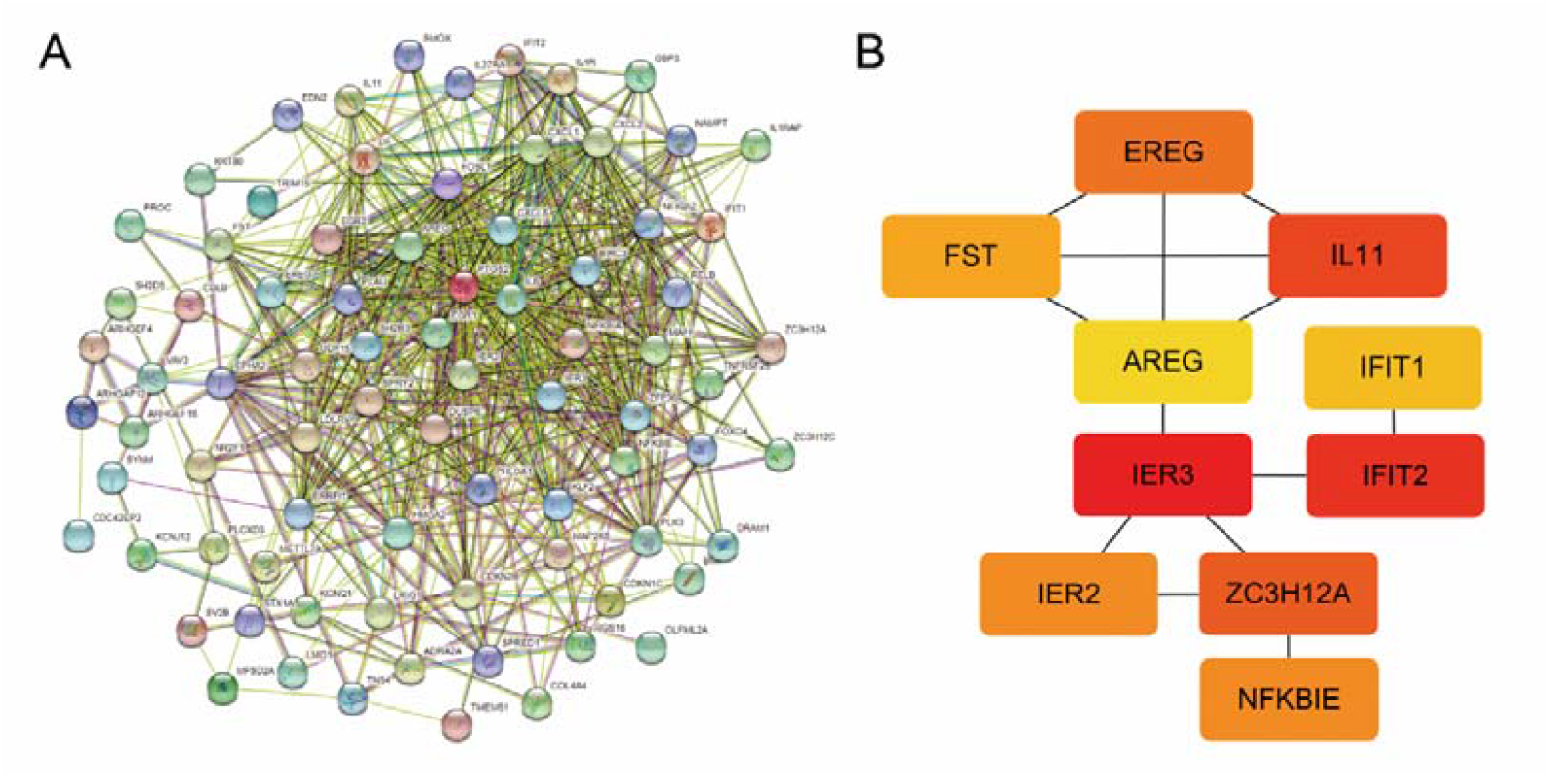
shows the PPI network for DEGs. A, PPI networks of significantly different genes after monkeypox infection of hela cells were generated using the STRING database; B, the top 10 hub genes were extracted by the Density of Maximum Neighborhood Component method in the cytohubba plug-in in Cytoscape.

### Monkeypox-related significantly different gene interactions and hub extraction

We constructed the interaction network of the significantly different genes using string database, and extracted the hub genes by DMNC method of SYTOHUBBA in cytoscape. The top 10 hub genes were IER3, IFIT2, IL11, ZC3H12A, Ereg, IER2, NFKBIE, FST,, iFIT1 and AREG(Table 2).

**Table 2.** The changes of 10 hub genes after monkeypox infection.

| Gene | logFC | P.Value |
| --- | --- | --- |
| IFIT2 | -1.56594 | 0.038802 |
| IFIT1 | -1.2736 | 0.01158 |
| IL11 | 2.777671 | 0.009122 |
| AREG | 2.840347 | 0.002257 |
| EREG | 2.689366 | 0.002555 |
| IER3 | 2.815185 | 0.004363 |
| IER2 | 1.0182 | 0.014747 |
| FST | 1.390166 | 0.008395 |
| ZC3H12A | 2.289584 | 0.022705 |
| NFKBIE | 1.11534 | 0.002333 |

### Verification of hub gene expression regulated by monkeypox virus

In order to verify the effect of monkeypox virus on the expression of the 10 hub genes, we obtained the data set of monkeypox-infected HELA in GEO (GSE36854 and GSE11234). As shown in Figure 6A-B, monkeypox significantly affected the expression of the 10 hub genes in dataset GSE11234, notably inhibiting the expression of IFIT1 and IFIT2. The expression data of IFIT1 and IFIT2 in the NCI-60 cell line drug database was downloaded from the CellMiner database, and the Pearson correlation coefficient with 792 FDA-approved drugs in clinical trials was analyzed. The results showed AP-26113 targeting IFIT1 and IFIT2 and Itraconazole targeting IFIT1.

**Fig. 6.**
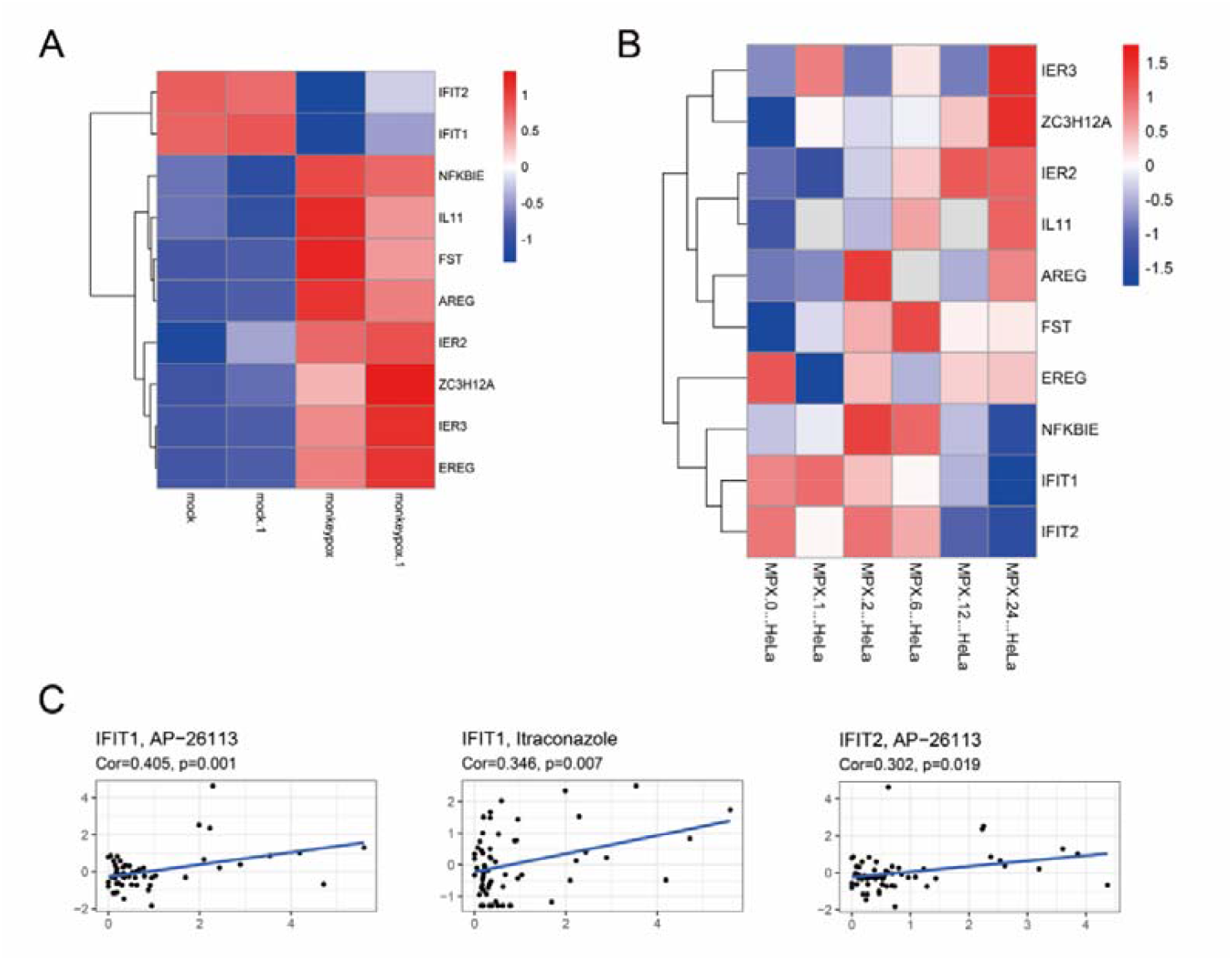
validation of the hub genes. A, heat map of the 10 hub genes in GSE36854 data set; B, heat map of the 10 hub genes in GSE11234 dataset; C, drug sensitivity evaluation targeting IFIT1 and IFIT2.

### Hub gene expression regulatory network

The gene expression is mainly regulated by transcription factors and miRNAs. In order to explore the relationship between hub genes and regulatory molecules, we used the network analyst online tool to deconstruct the core relationships of hub genes with key transcription factors and miRNAs. As shown in Figure 7A, the regulatory network of transcription factors with significantly different genes contains 16 nodes and 30 edges, with the major transcription factor IRF1, SIN3A, GLIS2, Smad5, ZFX, FOXJ2, and ATF1 in turn. As shown in Figure 7B, the regulatory network of miRNAs with significantly different genes contains 16 nodes and 30 edges. The key miRNAs are hsa-miR-16-5p, hsa-miR-21-3P, hsa-miR-520c-3p, hsa-miR-1343-3p, hsa-miR-203-3p and hsa-miR-335-5p. Given that IRF1 is the directly interacting transcription factors for both IFIT2 and IFIT1 in the regulatory network, we analyzed chip-seq data for IRF1 using the ENCODE database, which showed the presence of strong binding intensity of IRF1 on the chromosomes of IFIT family genes (Figure 7C).

**Fig. 7.**
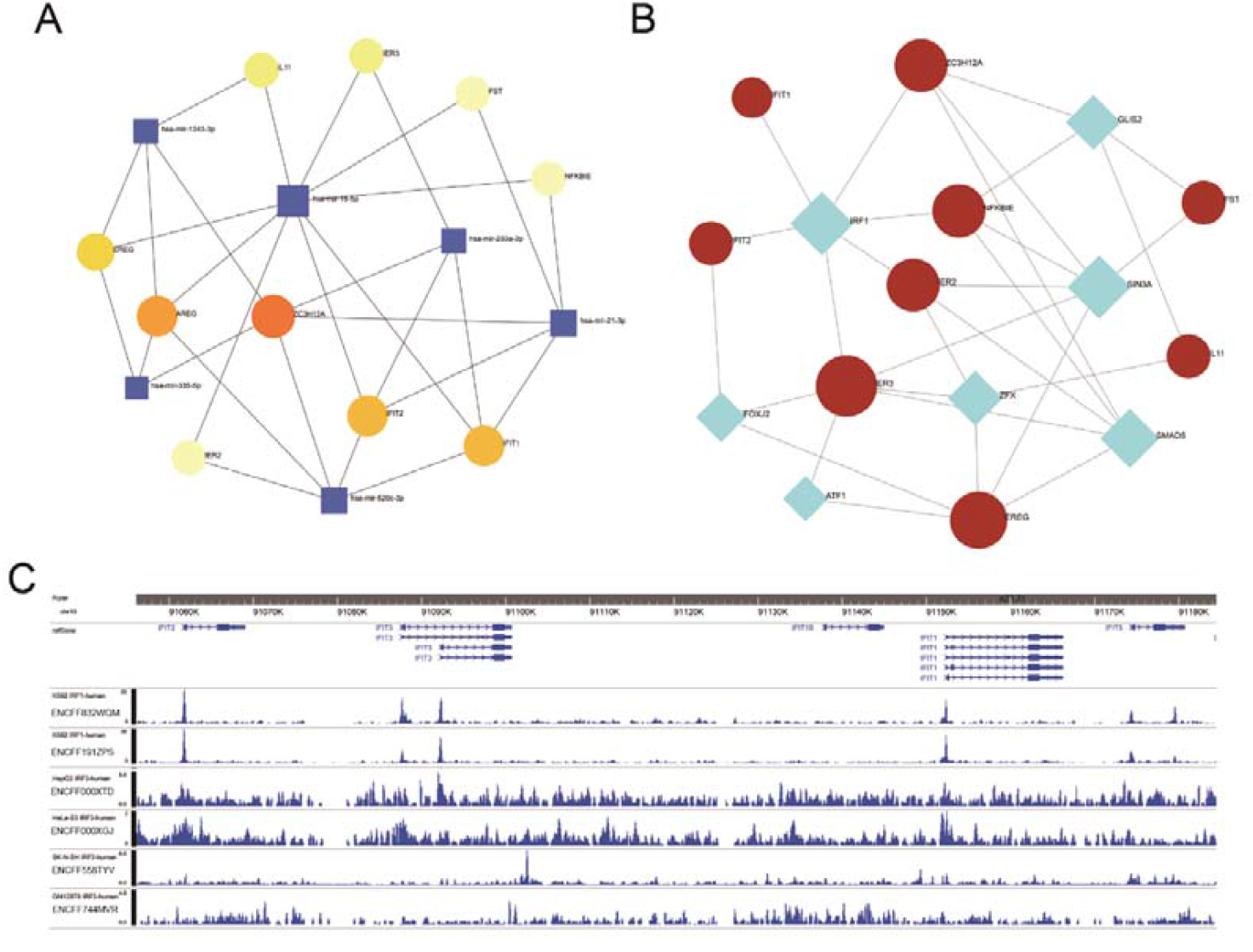
the expression regulatory network of hub genes. A, interaction network between hub genes and transcription factors based on NetworkAnalyst analysis; B, interaction network of hub genes with miRNAs based on the analysis of NetworkAnalyst; C, chip-seq sequencing data for IRF1 in the NCODE database on DNA binding strength of IFIT genes.

## Discussion

As monkeypox continues to spread around the world, it poses a major public health globally. For the fact that there were only a few sporadic cases of monkeypox in the past few decades, the research on the virus is limited to only case report, and only a few reports on the interaction between the virus and the host. To this effect, our current quandary in dealing with the massive spread of monkeypox is both crucial and so timely. Increasing the multi-dimensional research on monkeypox is something that both the scientific researchers should pay attention to also crucial for decision-makers. Of course, for researchers, clarifying the interaction between monkeypox and its host will be a prerequisite for dealing with monkeypox.

In view of the continuing difficulties in obtaining live monkeypox virus particles and the limited requirement for laboratory grade for monkeypox research, fortunately, there is sequencing data on monkeypox virus-infected cell models in the GEO data, which provides strong support for our study. We obtained two GEO sequencing datasets of monkeypox-infected Hela, GSE36854 and GSE11234, and screened 84 significantly differentially expressed genes (excluding histone genes) by analysis of differences and performed a series of analyses.

The viral early genes are extremely important for the viral infection, survival, replication and spread, also participate in the regulation of host immunity. We also analyzed the gene expression status of monkeypox virus itself, and screened out 26 possible early genes including D1L. The mechanism of how the early genes of monkeypox virus regulate the host and its life cycle is unclear, so it is urgent to study in this direction, which is also the focus of our next research. After obtaining the sequencing data of many monkeypox samples, we found that the 2022 epidemic monkeypox virus and the D1L gene reported in 2018 UK has multiple site mutations, given that D1L is monkeypox encoded Ankyrin repeat protein, we speculate that this may be the cause of monkeypox has become more adapted to human host and more adapted to human-to-human transmission.

Signal pathway analysis can effectively reflect the internal body in the real changes after external stimulation[27]. So by identifying the DEGs of monkeypox-infected Hela cells, we found that these DEGs were involved in the predecessor 10 KEGG signaling pathway, including TNF signaling pathway,C-type lectin receptor signaling pathway,NF-kappa B signaling pathway,Cytokine-cytokine receptor interaction,IL-17 signaling pathway,Th17 cell differentiation,Human T-cell leukemia virus infection, NOD-like receptor signaling pathway, Kaposi sarcoma-associated herpesvirus infection and Small cell lung cancer. These results suggest that monkeypox infection can activate the body’s strong immune response, produce an inflammatory response, and eventually lead to the body’s pathological changes.

People with monkeypox usually have high fever, headache, back pain, general malaise, cough, and swollen lymph nodes. Medical complications such as fever, rash, lymphadenopathy, pneumonia, encephalitis, eye-threatening keratitis, and subsequent bacterial infections may occur during the disease.[5]. Different diseases can be linked by their intrinsic genetic similarity[28]. We therefore performed genetic disease (GD) analysis to predict the association of Monkeypox DEGs with different diseases, and the results showed that diseases associated with monkeypox infection included Liver Cirrhosis,Myocardial Ischemia,Inflammation,Reperfusion Injury,Mammary Neoplasms,Hypertensive disease,Brain Ischemia,and Juvenile arthritis. This is consistent with the results of complications and sequelae of monkeypox. Additionally, my results show that monkeypox infection is associated with Schizophrenia and Mental depression, which supports the conclusion that monkeypox infection can lead to psychiatric disorders, such as anxiety, depression, and depression responses[29, 30]. Prostaglandin-endoperoxide synthase 2 (PTGS2), also known as cyclooxygenase 2 (COX-2), is the rate-limiting enzyme in the production of prostaglandins (PGs), plays an important role in various tumors and is also an important marker of iron death signaling pathway[31, 32]. PTGS2 plays an important role in the infection process of many viruses. Respiratory syncytial virus (RSV) and parafenfluenza virus infection lead to up-regulation of PTGS2 expression in airway bronchioles and bronchial epithelial cells and macrophages[33]. PTGS2 also promotes dengue and Sapovirus replication[34, 35]. All these indicate that PTGS2 can be used as a potential target for antiviral drug development. The role of PTGS2 in the process of monkeypox infection has not been reported, but given the degree of association between PTGS2 and many diseases, it is reasonable to speculate that monkeypox virus regulates the pathological process by regulating PTGS2. Of course, the current study of monkeypox is still infancy, the symptoms and sequelae caused by it still need long-term and in-depth follow up study.

The key gene (hub gene) plays a key role in the biological process. By extracting the hub gene from the protein-protein interaction network, we can effectively grasp the key molecules in the process of monkeypox infection. We extracted the hub genes of monkeypox-infected HELA cells including IER3,IFIT2,IL11,ZC3H12A,EREG, IER2,NFKBIE,FST,IFIT1 and AREG. The expression of the early response gene immediate early response 3 (IER3), formerly known as IEX-1, is induced by a great variety of stimuli, such as growth factors, cytokines, ionizing radiation, viral infection and other types of cellular stress[36]. IFIT genes are usually silent or expressed at very low constitutive levels. The induction of IFIT transcription is triggered by a number of stimuli, usually in the context of viral and bacterial infections[37]. IFIT is usually induced by type I and Type III interferons, with a much weaker induction of IFIT by Type II Interferons[38, 39]. RNA viruses such as respiratory syncytial virus, influenza viruses, West Nile Virus and vesicular stomatitis viruses are then exposed to TLRs, RLRs, and NLRS, resulting in large amounts of IFIT induction in the cell.

DNA viruses (e.g. Human herpesvirus 1 and Cytomegalovirus) that activate DNA sensors or cyclic GMP-AMP synthase [cGAS] also induce large amounts of IFIT expression[40-42]. IFIT1 and IFIT2 can inhibit mRNA translation initiation by binding to multi-subunit eukaryotic translation initiation factor 3(Eif3) and interfering with the assembly of the pre-initiation complex (consisting of the 40S ribosomal subunit, EIF3, eIF2/GTP/Met-tRNAi, and EIF4F)[43-48]. Specifically, IFIT2 inhibits the replication of West Nile Virus and Rabies in the central nervous system. In addition, IFIT2 protected mice from lesions caused by vesicular stomatitis virus and Sendai respiratory virus (SeV) [49-51]. Similarly, IFIT1 is an effector molecule that limits viral translation, and limits infection of viruses that lack RNA 2’-O methylation by binding to mRNAs that lack RNA 2’-o methylation[52-55]. Given the positive role of IFIT in antivirus, viruses have evolved multiple strategies to evade the antiviral capacity of IFIT[56, 57]. Different from other viruses, our results showed that monkeypox virus could significantly inhibit the expression of IFIT1 and IFIT2 in Hela cells. It may be that both of them have evolved some mechanism to regulate the expression of IFIT and escape the antiviral effect of IFIT.

The regulation of key genes by transcription factors and miRNAs after viral infection is very important to the pathological process. Then we analyzed the expression regulatory network of hub genes to find the transcriptional and post-transcriptional regulatory factors of hub genes. It also enables us to better understand the interaction of transcription factors and miRNAs with hub genes. We found that the main transcription factors regulating hub genes were IRF1, SIN3A, GLIS2, Smad5, ZFX, FOXJ2, and ATF1, and miRNAs were hsa-mir-16-5p, hsa-mir -21-3p, hsa-mir -520c-3p, hsa-mir -1343-3p, hsa-mir -203-3p, and hsa-mir-335-5p. Although the major players in viral-activated signaling pathways are IRF-3 or IRF-7, IRF-1 may also be a transcription factor associated with activation in some cells[58, 59]. IFIT2 and IFIT1 were shown to be potentially regulated by IRF1 from our results, which were validated in Chip-seq data for IRF1 in ENCODE data. Moreover, these miRNA functions are all virus-related, such as hsa-mir -520c-3p, which can be up-regulated by HBV and drive the invasion and migration of liver cancer cells [60] and hsa-mir-16-5p, which is an essential miRNA during infection with *E*.*histolytica* and is also significantly induced by HSV and HIV [61-63]. Hsa-mir -21-3p can promote IAV replication by inhibiting HDAC8[64]. There is no current report on monkeypox versus-related miRNAs, but given the important role of miRNAs throughout the course of viral infection, we have reasons to believe that miRNAs will be a hot spot in the next studies targeting monkeypox.

There is currently no effective treatment for monkeypox, and several antiviral drugs are thought to be effective against monkeypox. However, these drugs used to treat smallpox virus require scientific evaluation for the treatment of monkeypox[65]. Tecovirimat (TPOXX) is considered to be the most promising drug for monkeypox treatment. The mechanism of action of TPOXX is to target and inhibit the activity of VP37 protein of orthopoxvirus (a highly conserved gene encoded by all members of orthopoxvirus genus), and block their interaction with cellular Rab9 GTPase and Tip47, thereby preventing the formation of viral particles and inhibiting the transmission of the virus within an infected host[4, 66]. We performed drug sensitivity analyses through the CellMiner database and found that AP-26113 (Brigatinib), a tyrosine kinase receptor inhibitor and antitumor drug, significantly promoted the expression of IFIT1 and IFIT2, for the treatment of some forms of advanced non-small-cell lung carcinoma [67]. But there have been no reports of AP-26113 being used as an antiviral drug. In addition, Itraconazole, a broad-spectrum triazole fungicide, which is also widely used in the fight against influenza and HIV, significantly promotes IFIT2 expression[68, 69]. Because Itraconazole is also highly active against SARS-CoV-2 *in vitro*, it has also been applied in combination with other drugs to treat Covid-19 infections[70, 71]. There are no reports of Itraconazole against monkeypox, but based on its broad spectrum of antibacterial and antiviral effects, Itraconazole holds promise for the treatment of monkeypox infection in combination with other drugs. The above analysis is helpful for further experimental verification of the interaction between monkeypox virus and its host.

## Conclusion

Monkeypox virus significantly inhibited anti-viral gene IFIT1 and IFIT2 and reduced a variety of cell-intrinsic metabolic activities. AP-26113 and itraconazole promoting the expression of IFIT1 and IFIT2 may be used as new candidates for the treatment of monkeypox virus infection.

## Ethical Approval

N/A.

## Authors’ contributions

YT and GW designed this work. ZT and OB wrote the paper. ZT analyzed the data. YM, YM and XQ validated the analysis. All authors read and approved the final manuscript.

## Availability of data and materials

The data used and/or analyzed during the current study are available from the corresponding author on any reasonable request.

